# Automated refinement of metagenomic bins and estimation of binning success using *itBins*

**DOI:** 10.64898/2026.03.30.715291

**Authors:** Julian M. Künkel, Till L. V. Bornemann, Wei Xiu, Joern Starke, Tom L. Stach, André Rodrigues Soares, Jörg Schlötterer, Christin Seifert, Alexander J. Probst

## Abstract

Most prokaryotic genomes in public databases are genomes reconstructed from metagenomes, forming a compendium of multiple contiguous sequences (contigs) assembled from shotgun sequencing data. Binning algorithms for assigning contigs to metagenome assembled genomes (MAGs) are manifold and continuously improving in accuracy. However, binning errors, i.e. the incorrect assignment of a contig and coding sequence to a MAG, often propagate through various databases and confound taxonomic, metabolic and/or evolutionary analyses. Here we present itBins, a fully automated python-based software that enables ultra-fast refinement of metagenomic bins using a rule-based approach harnessing information from %GC content (%GC for brevity), coverage, and taxonomy of individual contigs. When applied to the low, medium, and high complexity data of the Critical Assessment of Metagenome Interpretation (CAMI I) challenge [1], itBins produced higher F_1_ scores (the harmonic mean of precision and recall) for all levels compared to other automated refinement tools, i.e., MDMcleaner and Rosella. Compared to manual refinement via uBin, itBins performed similarly well across all three complexity levels of the CAMI I dataset. With an average speed of 61 ms per bin, itBins is faster than all other refinement tools by at least three orders of magnitude when input data is accordingly available (%GC, coverage, and taxonomy), and was similarly fast when input data preparation was included in the processing time. Application to 64 real-world metagenomes from highly complex river mesocosms resulted in 259 medium-quality and 19 high-quality MAGs refined by itBins, while the other automated refinement tools failed in generating output at all or within 5000 hours of runtime. Finally, itBins also utilizes marker genes to determine the overall binning success for individual metagenomes, providing a crucial benchmark for the user to estimate the ecological relevance of their binned data. The herein introduced software itBins is broadly applicable to any type of metagenome data, integrates well with other software like DASTool, and enables swift and reliable refinement of genomes from metagenomes along with estimation of the overall binning success. itBins is distributed via EUPL 1.2 license and available at Codeberg (codeberg.org/JMK/itBins), GitHub (github.com/ProbstLab/itBins) and through Bioconda [2](bioconda.github.io/recipes/itbins/README.html).

## Background

Metagenomics as the major source of genomes from environmental samples has revolutionized our understanding of microbiology. It is based on the extraction of mixed-community DNA from an environmental sample followed by shotgun sequencing, for which a variety of different techniques exist. While read-based metagenomics focuses on the analysis of the short snippets of DNA (so-called reads) generated during sequencing, assembly-based metagenomics is based on the reconstruction of contiguous sequences (contigs) from multiple reads. Although losing data during the assembly process, the generation of contigs overcomes an inherent bias in short-read metagenomics, as reads only partially cover genes and thus taxonomic and functional assignments are hampered. Assembled metagenomes also enable the reconstruction of genomes from individual microbial populations, a technique also referred to as genome-resolved metagenomics. This results in so-called metagenome-assembled genomes (MAGs) [3, 4]. While this process loses even more data, it enables genome-centric analyses, such as phylogenomic profiling, metabolic potential analyses and network analyses of inter-microbe interactions or virus-host interactions. Genomes from isolates, in 2025, represent a minority in GTDB [5] compared to the estimated 99% of so far uncultivated prokaryotic microbes. While it would be preferable to acquire isolates of each species, this is currently infeasible, therefore MAGs are the foundation of multiple databases, aid other omics analyses such as proteomics, and enable vast evolutionary analyses across a major portion of the tree of life. For example, the study of the diversity of sulfate-reducing bacteria was substantially bolstered by the availability and reconstruction of MAGs [6], as was coining the concept of metabolic handoffs between microbes [7].

The reconstruction of genomes from metagenomes has led to the emergence of a variety of different software tools, referred to as binners (from genome binning, i.e. the assignment of contiguous sequences to one composite genome). While there is no universally best binner available across different environmental samples, consolidation software like DASTool [8] or MetaWRAP [9] unify the strengths of multiple binners applied to the same sample. Although these approaches can generate thousands of MAGs, the resulting composite genomes are often not of high quality [8] and many of them are not released into public databases or contain substantial binning errors that might propagate through databases, analyses, and scientific studies. Manual refinement of MAGs, which is possible via specifically designed tools included in the Anvi’o suite [10] and via uBin [11], has thus been frequently recommended, yet infrequently applied likely due to time limitations when processing bins. In similar fashion, automated refinement tools are also infrequently applied highlighting the necessity of a software for genome refinement that can be easily installed and applied.

We introduce itBins, an automated software for bacterial and archaeal MAG refinement which uses %GC, coverage and taxonomic information of individual contigs from a metagenome to evaluate the previously made assignment of contigs to a MAG. In addition, itBins uses marker genes across recovered MAGs compared to the corresponding metagenome to determine the overall success of a binning approach. This software is set out with the aim to automatically enhance the quality of genomes that are not manually refined and to counter error propagation in public databases by increasing reliability and reproducibility of metagenomic data.

## Methods

### The itBins algorithm

For the implementation of the itBins algorithm, python3 was selected as the programming language due to its widespread support and popularity in the scientific community, enabling researchers to engage in future extensions and development of the software. Dependencies were avoided where possible, with the exception of the python packages Pandas [12] and NumPy [13].

In easy terminology, itBins uses %GC, length, coverage (obtained, *e.g*., via mapping reads to the contigs), a taxonomy string representing the consensus taxonomy of all genes on the contig, and bacterial as well as archaeal single copy genes (SCGs) identified to be present or absent in each contig (*e.g*., obtained from searching predicted proteins on the contigs against a database). The SCGs used as default in itBins are those used in DASTool [8] and uBin [11, 14]. However, the source of this information is not static, as any source can be used. For example, usage of extensive single copy genes from software like CheckM v1 [15] can also be easily integrated, in cases of refining bins from specific lineages.

***Figure 1*** shows the conceptual workflow of the algorithm. It first loads the configuration file and the input data of one or multiple binned metagenomes, yet evaluates each bin individually. It then iteratively processes each candidate bin in the input, going through the refinement tasks, potentially multiple times. The refinement either stops when the last step or a preset stop task have been reached or an unrecoverable error has been encountered. The algorithm tasks can be configured individually.

**Figure 1.**
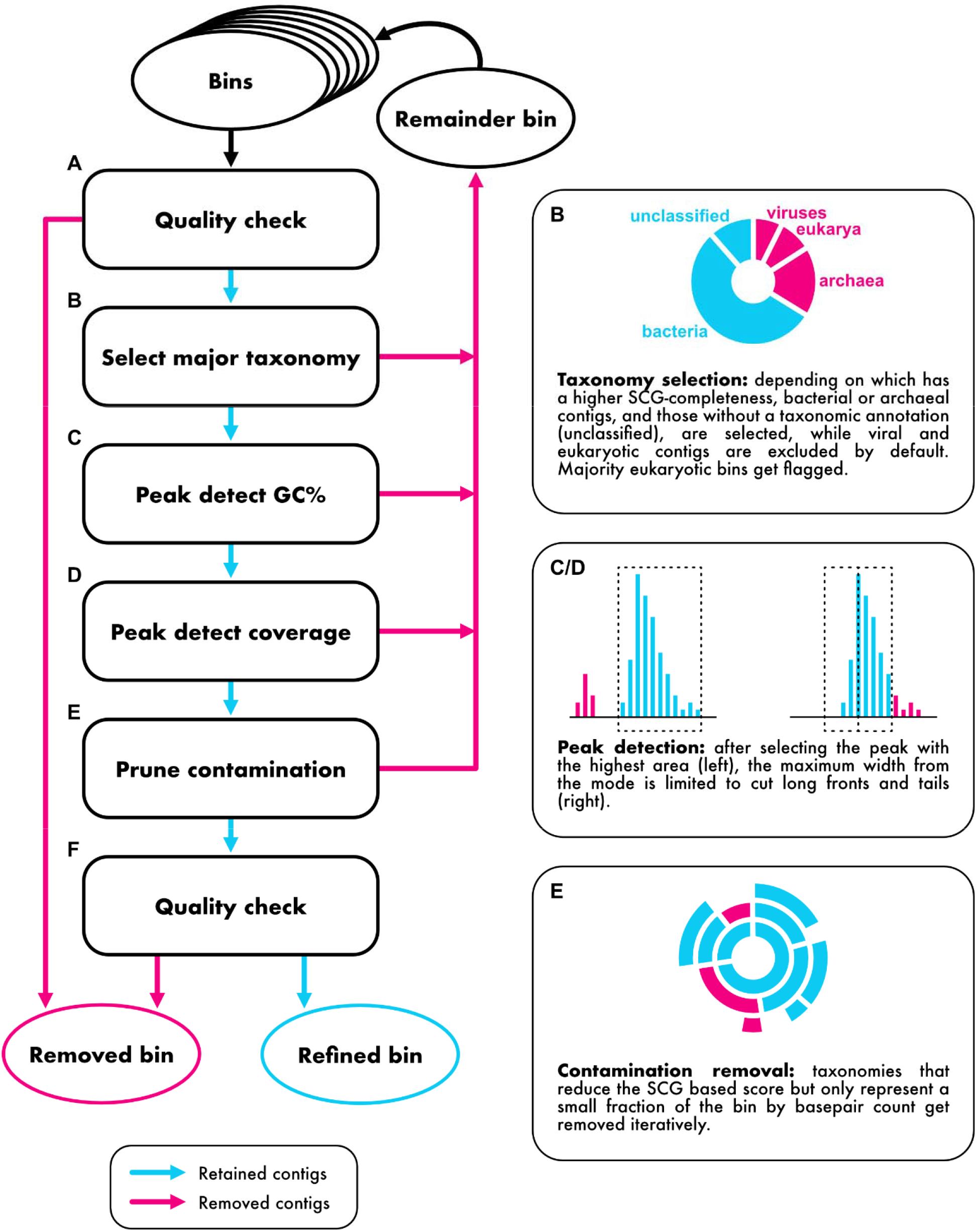
Conceptual workflow of the itBins algorithm to remove contamination from archaeal and bacterial bins.

By default, the eukaryotic fraction of the bin is calculated based on taxonomic information. If the bin crosses a definable threshold, it is flagged as a potential eukaryotic bin in the summary. These bins could be further refined with specific, user-provided marker gene sets, *e.g*. from BUSCO [16]. Afterwards, the archaeal and bacterial SCG completeness is calculated, to skip working on low completeness bins (< 70% completeness as default, yet definable, ***Figure 1 A***). Then, contigs labeled as viral or eukaryotic get removed from the bin, as well as either archaeal or bacterial contigs, based on which domain’s SCG completeness is higher (***Figure 1 B***). The refinement continues with contig exclusion based on its %GC. Specifically, after peak detection of the ideally uniformly distributed %GC of contigs in a bin, the peak with the highest area is selected. Then, if the edges of the peak are not at zero, the edges are extended until they hit the baseline excluding all contigs in the removed section. A check is then applied to limit overly wide peaks, by calculating a maximum width of the mode (***Figure 1 C***). The refinement based on coverage uses the same functions as for %GC based refinement, but in calculating the maximum peak width, it increases linearly with the coverage of the central peak to account for wider peaks at higher coverage (***Figure 1 D***). The taxonomy-based refinement removes contigs assigned a minority taxonomy (< 1% of the total length) that lower the overall bin score. The same scoring used in DASTool [8] is used here as well (***Figure 1 E***). The algorithm then calculates the metrics (completeness, contamination, score) for the bin, and finally excludes bins that do not meet the quality standards that were initially specified (default: completeness ≥70%, contamination ≤ 10%, ***Figure 1 F***). All specifications can be set in a config file supplied to the software.

Studies using metagenome-assembled genomes typically directly continue analysing the recovered MAGs to gain insights into the ecosystems, without evaluating how well these recovered MAGs actually represent the diversity of the environment they represent. This additional step of evaluating the representation of the metagenome by the recovered MAGs is implemented as the second module of itBins. The software analyzes single-copy gene distribution to estimate the overall number of binnable MAGs in a sample and compares those to the exact same genes represented in its output MAGs. These three marker genes included the bacterial *gyrA* and *rpS3* genes, and the archaeal *rpS3Ae* gene. For each of these genes, a rank abundance curve of the metagenome is computed. Contigs containing one or more of these genes are compared against bins and against the unbinned data for categorization of successfully binned or not successfully binned ranks in the rank abundance curves. In addition to the metrics for the whole dataset, a subset is analysed, formed by ordering the contigs in descending order of coverage and collecting the contigs that sum to 70% of the total coverage of ranked contigs. Contigs with a coverage < 7x are excluded from these estimations, as these are less likely to be assembled correctly [17]. We then report the result of this categorization in two ways: i) ‘absolute number of binned marker genes’, *e.g*., 312/503 and ii) ‘fraction of binned marker genes’, *e.g*., 0.62.

### Code availability, installation, and input data preparation

The code for itBins is publicly available via Codeberg (https://codeberg.org/JMK/itBins) and mirrored to GitHub (https://github.com/ProbstLab/itBins) under the EUPL 1.2 license. These resources also contain an installation guide for itBins.

The input data format for itBins is identical to those of DASTool and uBin. Ideally, these files are produced within a regular metagenome pipeline, yet their individual information and formatting are listed here. The software requires two input files: an overview file, in tab-separated-format, and a single-copy-gene-file, in comma-separated-format. The overview file lists length, %GC, coverage, taxonomy and bin assignment for each scaffold. The single-copy-gene-file lists the number of occurrences of each marker-gene for each contig. The files should be UTF-8 encoded. Descriptions of their fields are given in ***Table S2***, examples in ***Tables S3*** and ***S4***. Example input data is available on Codeberg and additionally at FigShare under https://doi.org/10.6084/m9.figshare.31157164.

### Evaluation of itBins and comparison to other refinement software

We compared itBins 0.8.3 to specific software that was designed for improving the quality of bins after automated binning. These specific software included MDMcleaner 0.8.7 [18], Rosella

0.5.4 [19], and uBin 0.9.20 [11], whereas the latter is a manual refinement software that uses identical parameters as itBins and thus serves as a general reference for the overall performance of itBins. For comparison of software performance (quality of output bins, run time) to the CAMI I challenge [1], the dataset contained a gold standard solution for each bin. For each complexity level of the CAMI I dataset, the S1 set was selected (for details please see original publication [1]); uBin data for CAMI I was retrieved from Bornemann et al. [11]. The F □ scores of the bins were calculated in relation to the gold standard bins, by first creating a list of all the gold standard IDs of the contigs found in the bin. From this a list of gold standard bin IDs was created, by recording which bin each contig was supposed to be part of. The gold standard bin IDs were sorted by the number of contigs linked to them. The bin with the greatest amount of links was assigned the corresponding gold standard bin. F □ scores were calculated from the number of missing contigs in the corresponding gold standard bin (false negatives), the number of true contigs assigned (true positives), and the number of falsely assigned contigs (false positives).

### Computational resources

All bin refinements reported herein were run on a server with 40 Intel(R) Xeon(R) E7-8891 v4 CPUs @ 2.80GHz, 1200GB DDR4 memory at 2400MT/s and Ubuntu 20.04.6 LTS “focal”. Network drives were accessible through a redundant 10G interface. The (shell built-in) ‘time’ command was used to determine the total time spent in user and kernel mode for the process.

## Results

### itBins successfully improves bins of the synthetic CAMI dataset

The first step of our evaluation of itBins was based on the three complexity levels of the CAMI I dataset, as these included gold standard bins which we used as the baseline for computing F □ scores from enumerated true positives, false positives and false negatives.

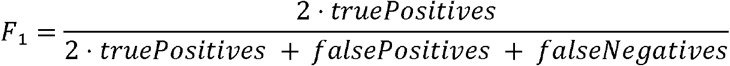

We compared itBins to MDMcleaner, Rosella, and uBin with regard to the quality of retrieved bins all computed from outputs of DASTool (please note that Rosella is also a binner and performs rebinning for refinement). The MDMcleaner output named “strictest”, “strict”, and “lenient” were considered for our analyses. “strictest” refers to only keeping contigs marked as “keep”, “strict” to keeping those marked as “keep” and “evaluate-low”, and “lenient” to keeping all that were not marked as “delete”.

Regarding the low complexity dataset, the bins refined with itBins had the highest median of F □ scores compared to those of all other refinement tools (***Figure 2***). However, we did not find significant differences in the F□ score distribution across all different datasets, reflecting the low complexity of the metagenomes resulting in binning with ease. By contrast, the medium complexity dataset resulted in the highest median F□ scores for uBin and itBins, respectively, yet only uBin was significantly different from the input dataset generated with DASTool. The performance of Rosella resulted in significantly lower F□ scores of bins compared to all other bin refinement tools (***Figure 2***). For the high complexity dataset, uBin and itBins significantly improved the F□ scores of bins generated with DASTool, while other refinement tools did not. Rosella again showed significantly lower performance compared to uBin, itBins, and MDMcleaner. At all levels of complexity, itBins performed worse, if provided with limited information only, skipping %GC, coverage or taxonomy based curation (***Figure S1***). Please note that MDMcleaner failed to refine 19 out of 30 bins for the low complexity, 29 out of 60 bins for the medium complexity, and 27 out of 75 bins for the high complexity datasets (***Figure S3***).

**Figure 2.**
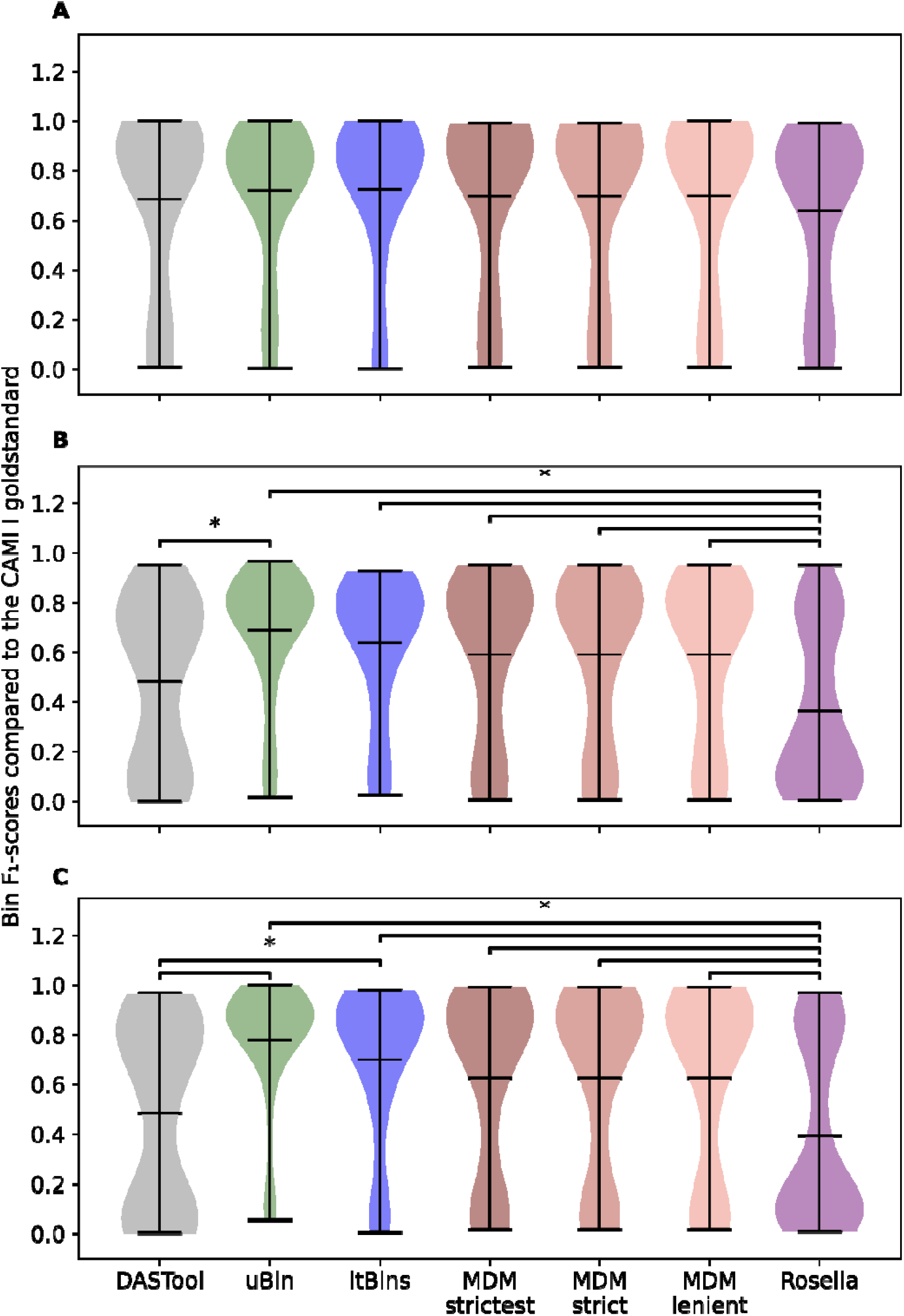
Violin plots of F□ scores compared to the CAMI I gold standard for the low (A), medium (B) and high complexity (C) sets. The F□ scores are shown on the Y-axis. The unrefined baseline after DASTool aggregation (DASTool), itBins automated refinement (itBins), manual refinement (uBin) and MDMcleaner automated refinement at different levels of strictness (MDM strictest, strict and lenient) and Rosella. Significant differences (marked *) were identified using Dunn’s test, and a significance level of 0.05.

### itBins outperforms other high-quality bin curators in run time

Besides the quality of the output, runtime of software for bin refinement is crucial as sequencing depth increases drastically [20] resulting in a constant expansion of genomic bins included in studies. Considering the CAMI I dataset and the individual run times across low, medium, and high complexity datasets, itBins outperformed all other automated refinement tools (MDMcleaner and Rosella) by four to five orders of magnitude (***Figure 3***) [18, 19]. Even when considering all the preprocessing, *i.e*., calculation of %GC, taxonomy, and coverage, which includes mapping of reads to contigs via Bowtie2 2.5.4 [21], itBins was substantially faster for low and medium complexity datasets. However, for the high complexity dataset, where itBins also outperformed the other two binners based on F□ scores (***Figure 2***) itBins remains slower than the other two curators when the time for preprocessing of data is taken into account. In sum, assuming that %GC, taxonomy, and coverage data are created for a new metagenome set out for binning, itBins outperforms both other automated curators, taking mere seconds to process the whole CAMI I dataset.

**Figure 3.**
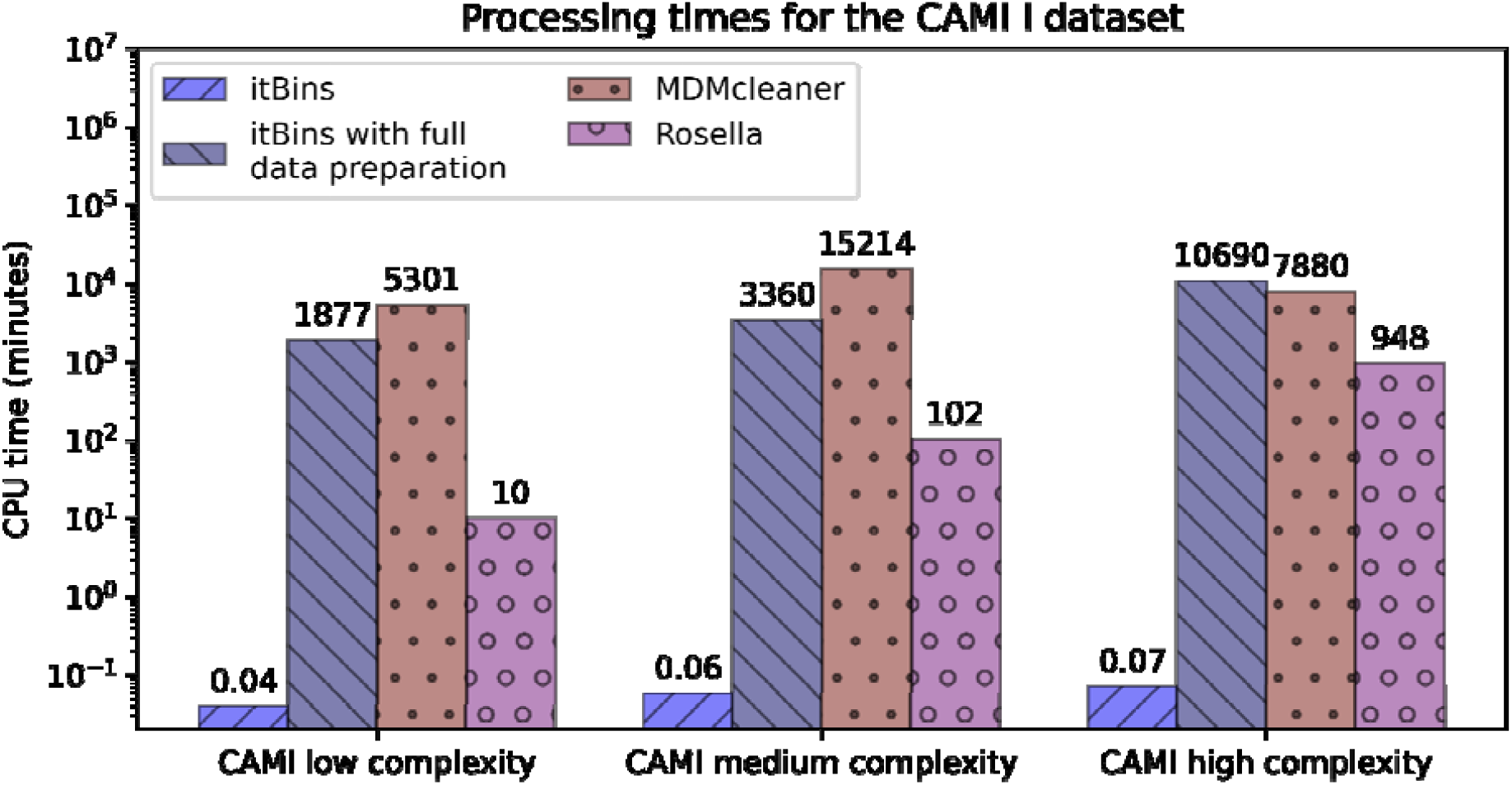
Processing times for itBins, itBins with full data preparation, MDMcleaner and Rosella, in minutes.

**Figure 4.**
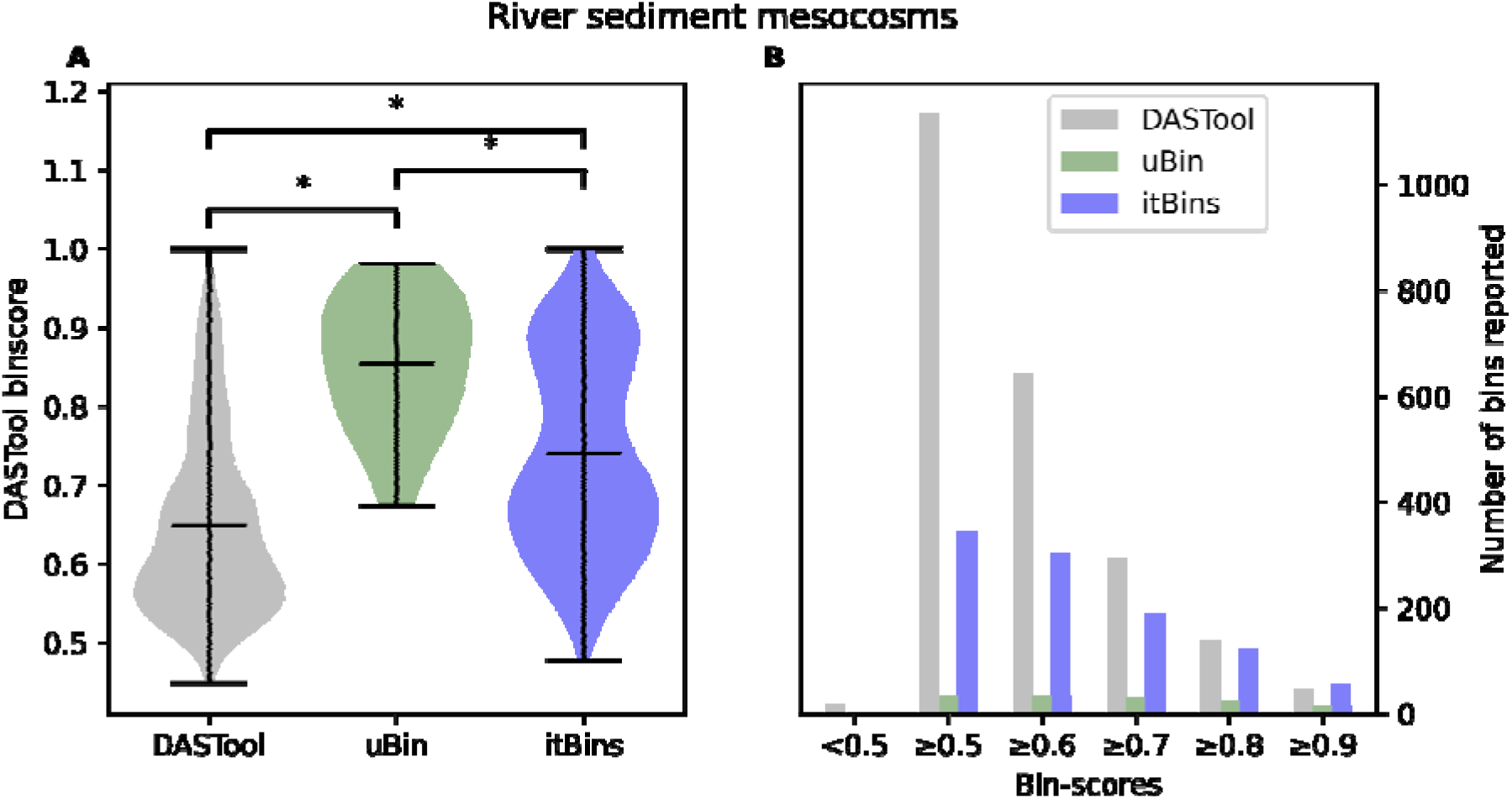
Application of itBins to real world datasets from a river mesocosm experiment spanning 64 metagenomes. A) Binscores of the river sediment mesocosm MAGs. Score-distributions are significantly different between the unrefined (DASTool), manually refined (uBin) and automatically refined (itBins) MAGs. B) Number of MAGs reported by binscore. DASTool reports a much higher number of MAGs compared to uBin and itBins for the lower thresholds. At a score greater 0.8, itBins reports a similar number of bins to DASTool, while removing or improving a large portion of the lower quality MAGs. At a score greater than 0.9, itBins reports a greater number of bins than DASTool. uBin consistently reports the lowest number of MAGs at scores greater than 0.5.

### itBins significantly improves metagenomic bins for real datasets

Data from river sediment mesocosm experiments [22] of high complexity (determined via Nonpareil [23]) was used to evaluate the performance of itBins on a real world, non-artificial dataset. The data encompassed 64 metagenomes available as unrefined bins and as bins manually refined using uBin. In lack of a gold-standard compared to CAMI I, *i.e*. ground truth contig-to-bin assignment, the binscore, as defined previously [8], was used as a metric of comparison. While refinement with itBins was a straightforward process, finishing 1525 bins after 17 minutes, MDMcleaner took 14,247 minutes, yet failed to create even a single MAG as output, due to segmentation faults, despite having previously processed many CAMI bins successfully. Rosella, on the other hand, was stopped by the authors as the software had not finished with any result after more than 300,000 CPU minutes for refining four of the 64 metagenomes. Considering how quickly Rosella refined the CAMI I datasets (***Figure 3***), this indicates poor scaling with the input size.

Both uBin and itBins managed to significantly improve upon DASTool in terms of binscore. Further, uBin achieved a significantly better result than itBins. Both uBin and itBins improve on the minimum score, with the highest score reported by itBins being higher than the highest score reported by uBin, but the lowest score reported by uBin was higher than the lowest score reported by itBins. Similar to the results for the CAMI I dataset, itBins performed worse when skipping %GC, coverage or taxonomy based curation (***Figure S2***).

### Estimating binning success with itBins

ItBins provides a success estimate for the user regarding the recovery of bins from the assembled metagenome. For doing so, itBins utilizes single copy genes for bacteria (ribosomal protein S3 (*rpS3*) and gyrase subunit A (*gyrA*)) and archaea (ribosomal protein S3Ae, rpsS3Ae). However, some genomes that are of low abundance in the sequencing dataset often do not result in high quality bins due to a lack of data; hence, itBins reports the overall success of considering all detectable genomes based on the abovementioned markers along with a revised dataset that only considers the top 70% of the marker genes based on coverage retrieved from read mapping. Applying these estimates to the CAMI I dataset revealed success rates between 27.0% and 74.2% recovery efficiency across the low, medium, and high complexity dataset (***Figure 5, Table S1***). Interestingly, the two different marker genes for bacteria performed equally well, although they can result in differing community profiles of natural samples [24].

**Figure 5.**
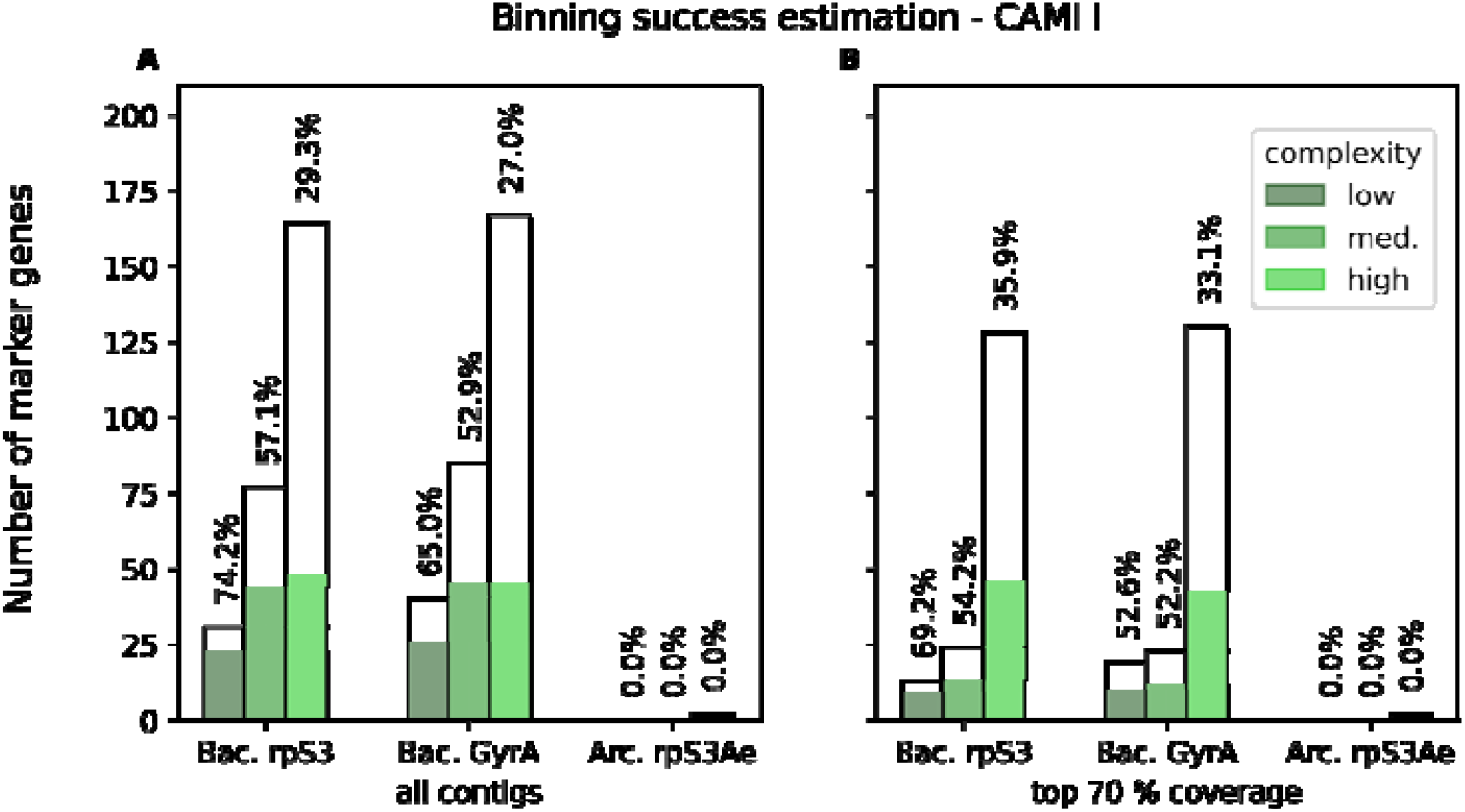
Binning success estimation for the CAMI I datasets via itBins, showing the binned (green) vs. unbinned marker genes (white). Both overall (A) and when only considering the contigs making up the top 70% of total coverage (B), the highest fractions of binned marker genes is found for the low complexity set, the lowest for the high complexity set.

Application of the binning success estimation to river sediment mesocosms of high complexity (see above) demonstrated that only 730 out of 16,123 (4.5%) of the bacterial *gyrA* genes were binned, with 475 of 9,985 (4.8%) in the top 70% based on coverage. Regarding the bacterial single marker gene *rpS3*, 877 of 11,722 (7.5%) were overall binned with 683 of 8,121 (8.4%) in the top 70% coverage. One of two archaeal *rpS3Ae* genes was successfully recovered in a MAG, falling in the top 70% (***Figure S4***). This dataset is highly diverse, likely shaped by strain heterogeneity [22] and more complex than even the CAMI I high complexity set. In sum, a large fraction of the community of the river sediment mesocosm study remains unbinned, with a higher proportion of the major players (represented in the top-70%-coverage metric) being recovered as genomes, leaving the rare biosphere mostly enigmatic regarding genomic resolution.

While classically the metric of binning success would be the percentage of reads mapping to MAGs, this marker gene based method is quickly achieved without another time consuming mapping, representing a low-to-no-cost alternative in terms of computing resources.

## Discussion

### Research areas that urgently necessitate bin refinement

Improvements in data processing and low requirements regarding knowledge in bioinformatics have made genome-resolved metagenomics accessible to most environmental microbiologists. It was software like Anvi’o [10] that enabled this progress, particularly due to its interactive visualization tools that made the software user friendly. Integrated in Anvi’o [10], manual refinement (anvi-refine) of metagenomic bins enables refinement of genomes similar to uBin [11]. However, there are only few tools available in the literature that enable users to automatically refine bins without time-consuming manual tinkering with the contig assignments.

Although Anantharaman et al. refined more than 2k metagenomic bins in their study [25], there are numerous studies whose size simply makes manual refinement of bins nearly impossible. For example, the recently published catalogue of about 2M metagenomic bins [26] would necessitate more than 50 years of manual refinement assuming only spending 3 min per bin for 40 hours per week. There are also much smaller studies from various fields [27–30] but particularly in the field of human microbiome research [31–35], that could substantially benefit from bin refinement. These also include a study led by Mahnert et al. [31] describing a catalog of more than 1k archeal genomes from the human microbiome which could have been manually refined as the dataset is much smaller than for Anantharaman et al. [25]. Yet, the human microbiome is usually less complex than soil or sediment microbiomes, and thus often does not benefit that much from bin refinement, as we showed for our CAMI I low complexity dataset (***Figure 2***).

In sum, we generally recommend automated refinement of bins as it usually does not make the data worse for low complexity datasets, yet highly commend the application to medium and high complexity datasets as it substantially improves the quality of many bins. This is also highlighted by the fact that post-refinement with itBins generated more high-quality bins for the river sediment samples analyzed herein than the regular DASTool output, i.e., many of the lower quality bins could be substantially improved. General application of bin refinement to genome-resolved metagenomics studies would generate more accurate data, make science more reliable, and avoid error propagation in public databases.

### User experience during application of bin refinement tools

Comparative analysis of refinement tools with data for the CAMI I dataset clearly demonstrated that itBins outperforms all other automated tools tested herein in speed and quality. However, apart from the sensitivity and efficiency of a new bioinformatics software, the user experience from installation to interpretation of results is equally important. When applying the different automated refinement tools in this study, we encountered some problems during the installation of MDMcleaner and the download of the respectively required databases. Only after modifying the source code, MDMcleaner could be installed and successfully built its databases. These modifications were related to the code checking the versions of installed dependencies, GTDB paths and checksum comparisons, but not to any parts of the code responsible for the refinement of metagenomes (the program is therefore assumed to be functioning as intended, versions/checksums were confirmed manually). During the application of MDMcleaner, the refinement of some CAMI I genomes failed, with inconclusive error messages pointing to problems with multiprocessing, which persisted even in single-thread mode. During refinement of the river sediment datasets, MDMcleaner crashed upon hitting segmentation faults. The program assigned one of four designations to the contigs, ‘keep’, ‘evaluate-low’ (low probability of containment), ‘evaluate-high’ (high probability of containment) and ‘drop’. Just using the contigs designated ‘keep’ is considered the strictest refinement mode, using ‘keep’- and ‘evaluate-low’-contigs is referred to as strict mode. Including all contigs but the ones designated ‘drop’ is referred to as lenient mode. Contigs designated ‘drop’ are never included. Yet, the MDMcleaner output is easy to understand and well structured, referencing the original bin names.

Rosella was able to perform automated refinement, despite some inaccuracies in its usage documentation, due to informative error messages. Rosella also renamed all modified bins, complicating comparisons and analyses. While Rosella was able to refine the small CAMI I dataset in quite competitive time, the process had to be terminated after 5,000 hours for the river sediment mesocosm dataset, indicating a poor scaling of its algorithms.

Building on these experiences, itBins provides a more streamlined user experience, with minimal dependencies and a simple install with conda. The software itself does not require the download or building of databases. The fast refinement with itBins, and its ability to reuse prior work (*e.g*., mapping or BLASTs [36]), make it an attractive add-on for metagenomics workflows. While itBins does not require its input to be in a form that would chain it to any specific software, it does require all the necessary metrics to perform its three core refinement tasks based on %GC, coverage and taxonomy, as shown by the degraded performance when omitting any of these. It flags bins that it cannot refine successfully or suspects to be eukaryotic; removed scaffolds are reported, and the software provides a summary with reasons for the flag, providing a transparent and explainable system. At the same time, the system is also tunable, by adjusting the refinement tasks’ parameters or providing custom marker gene sets, *e.g*., from CheckM [15]. itBins’ runtime scales linearly with the number of bins, and may be parallelised over them in the future, as each bin can be refined independently. Only input acquisition and binning success estimation as well as output aggregation and formatting will parallelise poorly, yet make up only a small proportion of runtime for large datasets. Collaborations with another lab on the refinement of 4075 metagenomic bins [37] ensured the robustness of the software, from installation over handling to interpretation of the results.

### Considerations for future refinement tools

While the basic principle and the used data behind itBins is quite simple, i.e., a rule-based approach utilizing %GC, coverage, and taxonomy, future binning tools should harness machine-learning-based approaches for bin refinement, including the use of large/genomic language models. Another layer of information besides those used by itBins, is the assembly graph generated during the metagenome assembly. Unfortunately, the software BinSPreader [38], which works off assembly graphs produced during metagenome assembly, could not be tested on the CAMI I dataset as no such graph is available for this dataset. Moreover, improvement and integration of approaches for chimerism should be considered in bin refinement. For example, GUNC reports bins with chimerism, yet the software does not disclose the genomic position of the chimerism thus not enabling refinement [39]. It also remains elusive, how well certain bin refiners perform on novel phylum-level lineages where taxonomy information is sparse or itself confounding. These are essential key facts to be considered for the design and implementation of future refinement tools.

With respect to itBins, several improvements could be made in the future regarding the data preparation workflow, especially the speed of mapping and taxonomic assignments, could greatly improve the user experience of itBins. We also envision that by incorporating an analysis of k-mer frequencies of lengths common in assembly, the %GC-refinement task may be improved, similarly, differential coverage refinement might improve the coverage-refinement task. itBins might also benefit from referencing the number of contigs in the bin when adjudicating peakwidth, as a more fragmented genome will naturally have a broader %GC-distribution. Similarly, longer contigs will have a %GC closer to the mode of the distribution, and thus long contigs with a difference in %GC to the bin average are more likely to be misassigned than shorter ones. In similar fashion, taxonomy-based refinement in itBins so far uses a singular taxonomic assignment per contig. By utilising the assignments of all ORFs on one contig as information input for itBins, a more holistic view of the taxonomy of a bin might enable more robust refinement decisions. Lastly, the bin score is calculated based on a set of universal bacterial or archaeal marker genes, with the possibility to use a provided, different set for each. More accurate scores might be possible, if marker gene sets are created at a taxonomic rank lower than domain. While itBins has room for improvement, it is also structured in a way that facilitates easy, maintainable improvement. For example, users can easily decide which tool they use for input generation, may it be a quick taxonomic profiling via one of the bbtools [40], quickclade (https://bbmap.org/tools/quickclade) or kraken2 [41] or importing coverage values for contigs that are generated via a much faster software like coverM [42].

## Conclusion

Bin refinement is important to correct errors in binners and bin aggregators, yet expensive, time-consuming and requires experience. The automated refinement tool itBins provides consistent, ultra-fast refinement, faster than existing tools even for large datasets. It is faster and more accurate than other tested automated bin refiners and produces similar results as those via manual refinement. itBins also estimates the success of the binning effort to provide a measure of the applied tools to the user and guide study design. Lastly, itBins is built modularly and ready to be improved or tuned when it comes to input data, while at the same time being flexible for integration into existing workflows of metagenomics labs around the globe.

## List of abbreviations

BLAST: Basic Local Alignment Search Tool
CAMI: Critical Assessment of Metagenome Interpretation
Contig: Contiguous sequence
CPU: Central Processing Unit
CRC: Collaborative Research Center
DASTool: Dereplication, Aggregation and Scoring Tool
DFG: German Research Foundation (Deutsche Forschungsgemeinschaft)
DNA: DeoxyriboNucleic Acid
EUPL: European Union Public License
GMEC: State Key Laboratory of Geomicrobiology and Environmental Changes
GTDB: Genome Taxonomy DataBase
GUNC: Genome UNClutterer
*gyrA*: bacterial DNA Gyrase subunit A gene
ID: IDentifier
MAG: Metagenome Assembled Genome
MDMcleaner: Microbial Dark Matter cleaner
ORF: Open Reading Frame
*rpS3*: bacterial ribosomal protein S3 gene
*rpS3Ae*: archaeal ribosomal protein S3Ae gene
SCG: Single Copy Gene
UTF-8: Unicode transformation format, 8-bit
ZMB: Center for Medical Biotechnology (Zentrum für Medizinische Biotechnologie)
ZWU: Center of Water and Environmental Research (Zentrum für Wasser- und Umweltforschung)

## Declarations

No declarations to make.

## Availability of data and materials

itBins is available via conda through the Bioconda channel. Metagenomic data presented in this study was re-used in accordance to Hug et al. 2025 [43]. The CAMI I dataset is available at https://doi.org/10.5524/100344, the unrefined MAG dataset available at https://doi.org/10.6084/m9.figshare.31157164, while river sediment microbiome data [22] is available via BioProject PRJNA1085635 with the redundant and unrefined MAG dataset available at https://doi.org/10.6084/m9.figshare.31796194.

## Authors’ contributions

JMK: Software design and implementation, software testing, data analysis, visualization, wrote first draft of manuscript, manuscript revision

TLVB: Software design and implementation, supervision of software design, manuscript revision WX: Software testing, manuscript revision

JSt: Software testing, manuscript revision TLS: Provided data, manuscript revision

ARS: Tested user experience, software design, manuscript revision JSc: Supervision of software design, manuscript revision

CS: Supervision of software design, manuscript revision

AJP: Designed project, software design, provided supervision, visualization, wrote first draft, manuscript revision

## Acknowledgements

We thank Ken Dreger for outstanding system administration. We thank Brian Bushnell for discussions. We thank Johannes Köster for help with the distribution of itBins. We also thank Till Bornemann, Sarah Esser, Carrie Moore, Jule Nuy, Julia Plewka, Alexander Probst, Manan Shah, Sophie Anja Simon, André Soares, Tom Lennard Stach, Jörn Starke and Katharina Sures for providing insights into their manual bin curation processes.

