## Supplementary Information for "Automated refinement of metagenomic bins and estimation of binning success using *itBins*"

Content:

1. Supplementary Figures 1-4
2. Supplementary Tables 1-4

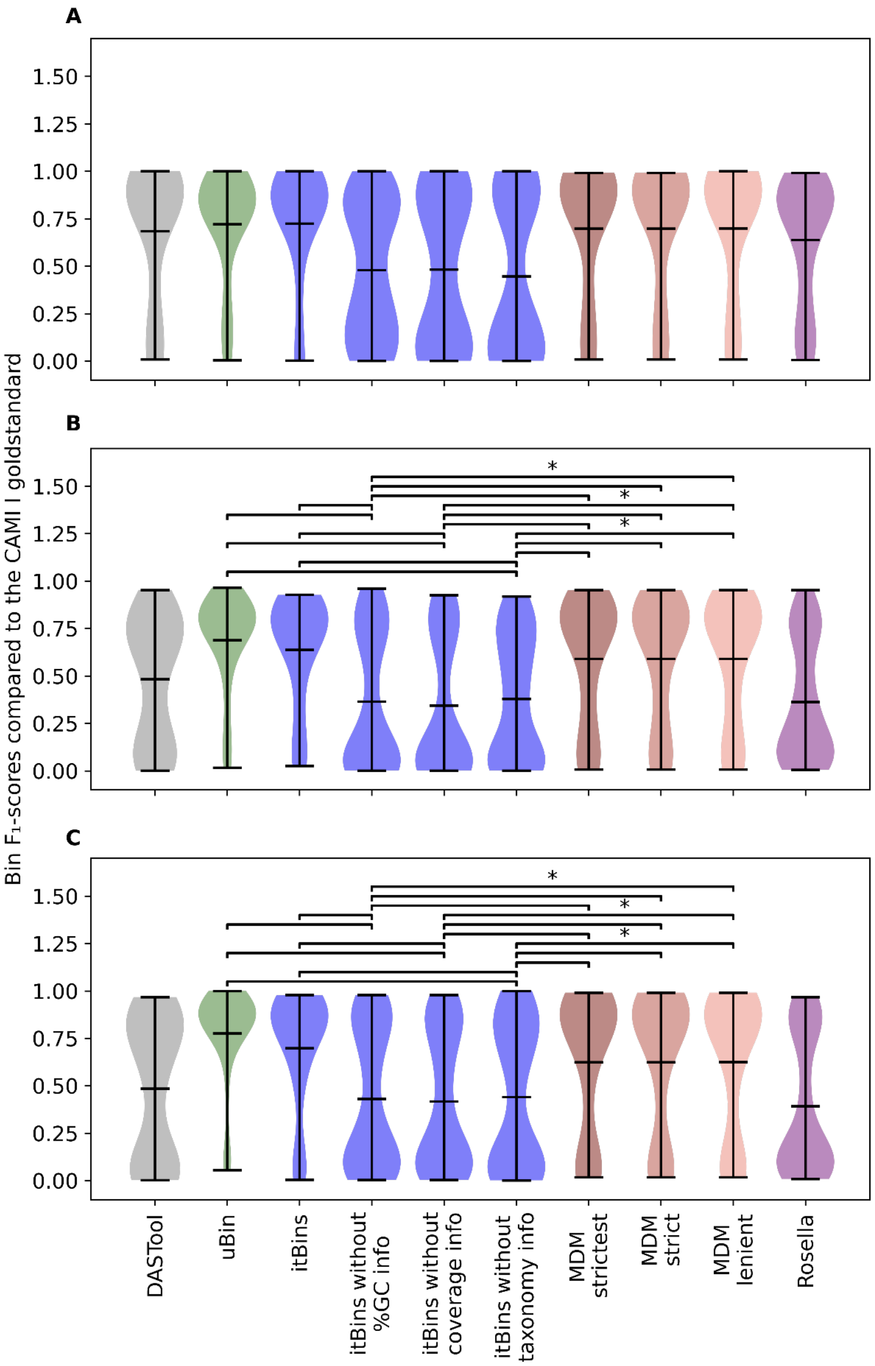


*Figure S1: Violin plots of $F_1$ scores compared to the CAMI I gold standard for the low (A), medium (B) and high complexity (C) sets. The $F_1$ scores are shown on the Y-axis. The unrefined baseline after DASTool aggregation (DASTool), itBins automated refinement (itBins), manual refinement (uBin) and MDMcleaner automated refinement at different levels of strictness (MDM strictest, strict and lenient) and Rosella. Significant differences were identified using Dunn's test, and a significance level of 0.05, and are only shown for itBins with limited input.*

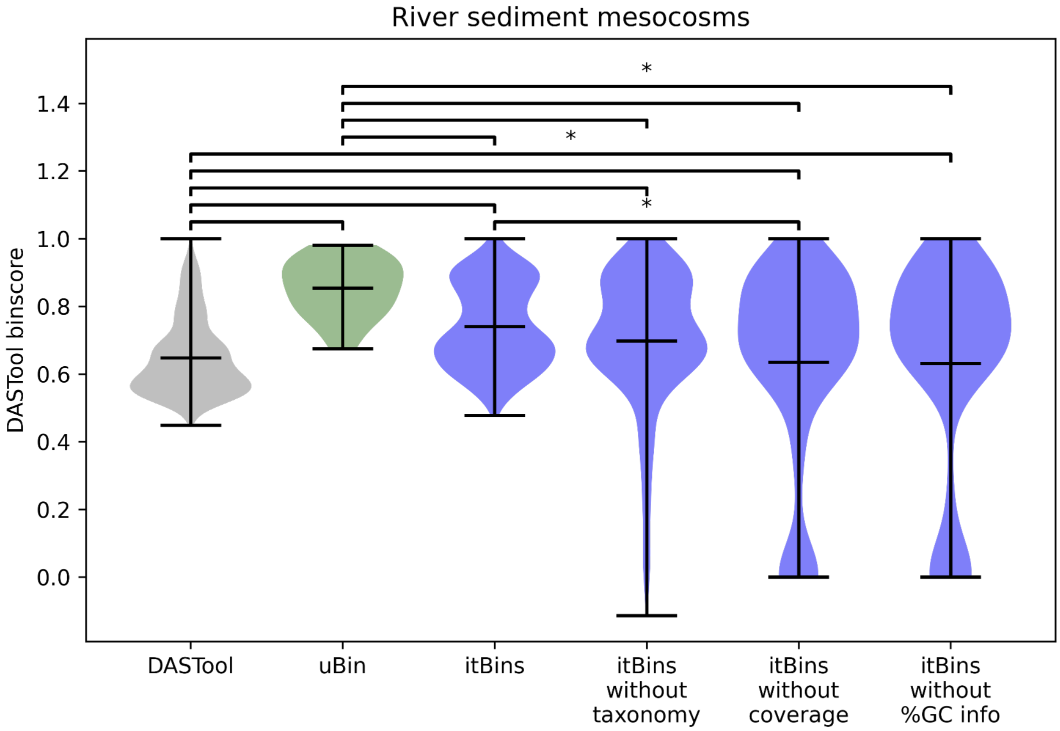


*Figure S2: Binscores of the river sediment mesocosm MAGs. In addition to DASTool, uBin and itBins, distributions are shown for itBins with limited input. Significant differences were identified using Dunn's test, and a significance level of 0.05.*

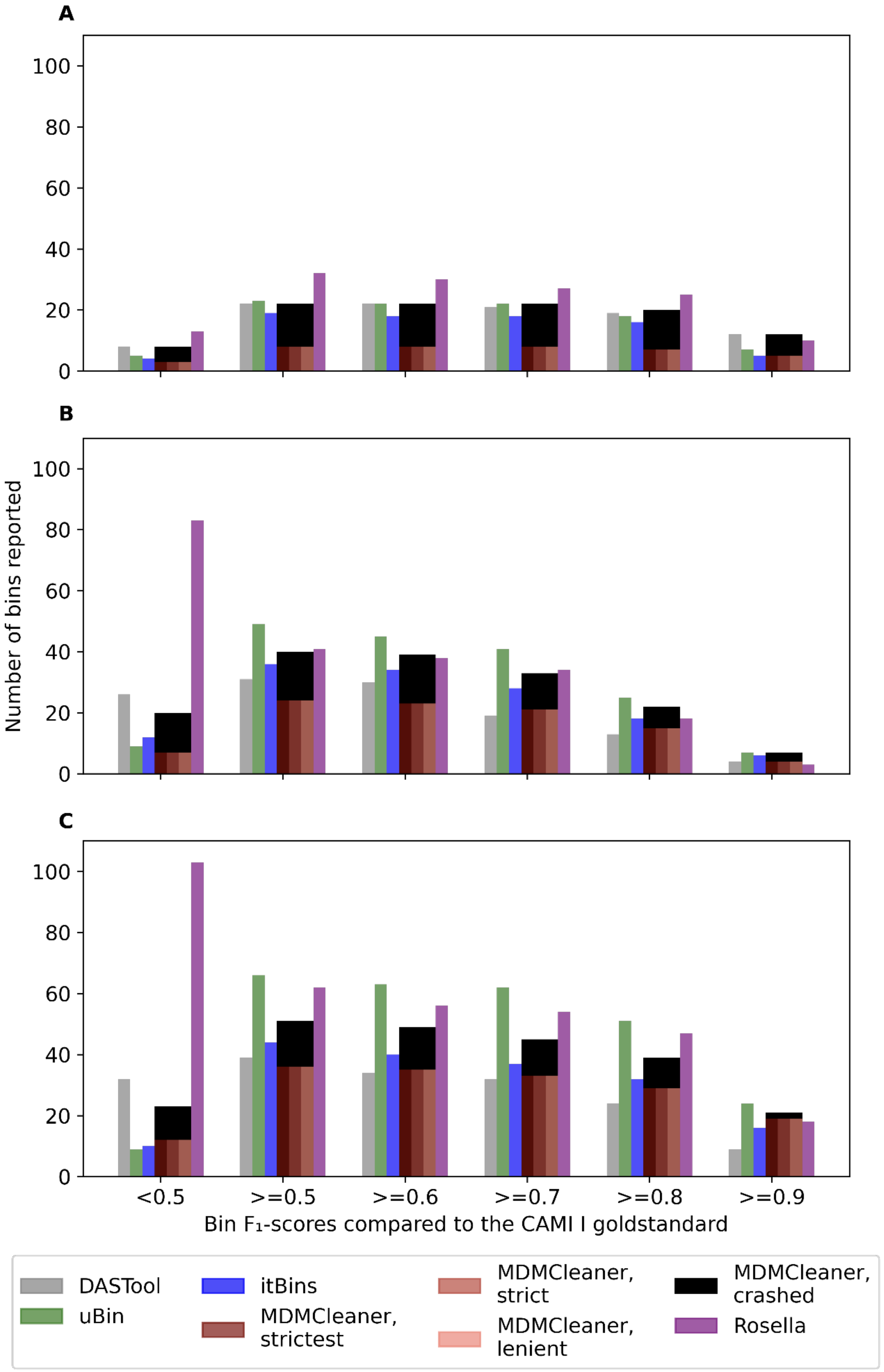


*Figure S3: Number of reported bins for DASTool, uBin, itBins, MDMcleaner and Rosella at $F_1$ score thresholds between 0.5 and 0.9, for the low (A), medium (B) and high complexity (C) sets.*

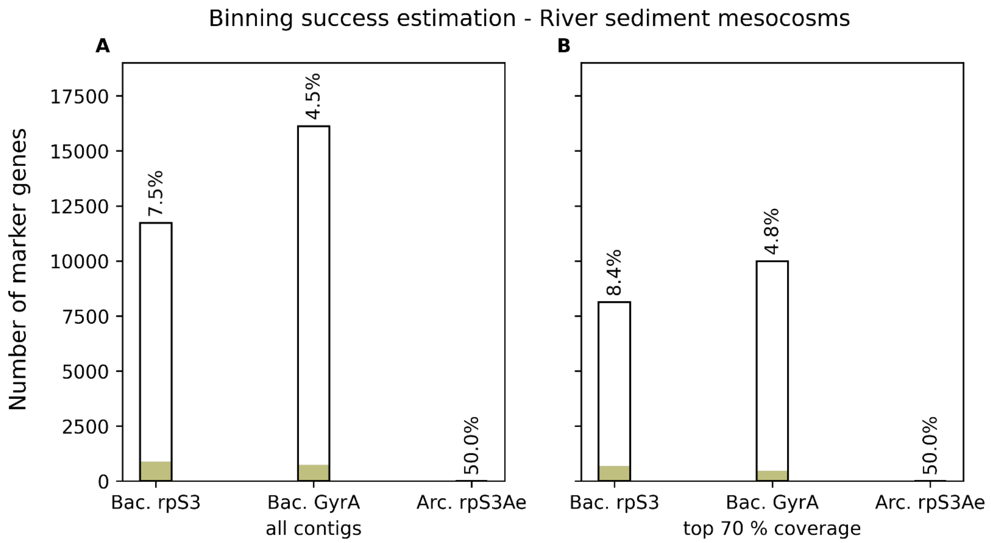


*Figure S4: Binning success estimation for the river sediment mesocosms, considering all contigs (A) and only the contigs making up the top 70 % of total coverage (B).*

Table S1: Binning success estimation for the CAMI I dataset

| CAMI set | Bac. *rpS3* overall | Bac. *rpS3* top 70 % coverage | Bac. *gyrA* overall | Bac. *gyrA* top 70 % coverage | Arc. *rpS3Ae* overall | Arc. *rpS3Ae* top 70 % coverage |
|---|---|---|---|---|---|---|
| low complexity | 23 of 31 (74.2 %) | 9 of 13 (69.2 %) | 26 of 40 (65.0 %) | 10 of 19 (52.6 %) | 0 of 0 (0.0 %) | 0 of 0 (0.0 %) |
| medium complexity | 44 of 77 (57.1 %) | 13 of 24 (54.2 %) | 45 of 85 (52.9 %) | 12 of 23 (52.2 %) | 0 of 0 (0.0 %) | 0 of 0 (0.0 %) |
| high complexity | 48 of 164 (29.3 %) | 46 of 128 (35.9 %) | 45 of 167 (27.0 %) | 43 of 130 (33.1 %) | 0 of 2 (0.0 %) | 0 of 2 (0.0 %) |

Table S2: Descriptions of the input data fields

| Field | Contents | Comment |
| --- | --- | --- |
| scaffold identifier | String representable as 256 bytes | Usually 256 characters, fewer in case of special characters |
| Bin identifier | String representable as 256 bytes | Usually 256 characters, fewer in case of special characters |
| Marker gene identifier | String representable as 256 bytes | Usually 256 characters, fewer in case of special characters |
| Taxonomic designation | Up to 6 semicolon-separated strings, each representable as 256 bytes | Per taxonomic rank, usually 256 characters, fewer in case of special characters. |
| %GC | Numeric string representable by an unsigned 16-bit integer with a scaling factor of 0.1 (0.0-100.0) | Usually two digits, followed by a dot, followed by a single digit |
| coverage | Numeric string representable by an unsigned 64-bit integer with a scaling factor of 0.1 (0.0-1.8*10^18) | Usually two to four digits, followed by a dot, followed by a single digit |
| length | Numeric string representable by an unsigned 64-bit integer (0-18*10^18) | Usually four to seven digits |
| Number of marker gene occurences | Numeric string representable by an unsigned 8-bit integer (0-255) | Usually a single digit. In the unlikely case of a contig containing more that 255 copies of the same marker gene, setting it to 255 should not skew results much |

Table S3: Overview file example

| scaffold | length | gc | coverage | taxonomy | bin |
|---|---|---|---|---|---|
| exampleContig_001 | 1053 | 55.3 | 9.1 | bacteria;examplePhylum; | exampleBin |

Table S4: Single-copy-gene file example

| scaffolds | b_example_geneA | b_example_geneB | b_example_geneC | … |
|---|---|---|---|---|
| exampleContig_001 | 0 | 1 | 0 | … |
